# West Nile virus and Usutu virus in wild birds from Rescue Centers, a post-mortem monitoring study from Central Italy

**DOI:** 10.1101/2022.07.19.500416

**Authors:** Giuseppe Giglia, Giulia Mencattelli, Elvio Lepri, Gianfilippo Agliani, Marco Gobbi, Andrea Gröne, Judith M. A. van den Brand, Giovanni Savini, Maria Teresa Mandara

**Affiliations:** Department of Veterinary Medicine, University of Perugia, 06126 Perugia, Italy; Division of Pathology, Department of Biomolecular Health Sciences, Faculty of Veterinary Medicine, Utrecht University, 3584 CL Utrecht, The Netherlands; OIE National Reference Center for West Nile Disease, Istituto Zooprofilattico Sperimentale dell’Abruzzo e Molise “G. Caporale”, 64100 Teramo, Italy; Istituto Zooprofilattico Sperimentale dell’Umbria e delle Marche “T. Rosati”, 06126 Perugia, Italy; Dutch Wildlife Health Centre, Utrecht University, 3584 CL Utrecht, the Netherlands

**Keywords:** Arboviruses, West Nile, Usutu, Monitoring, Wild Birds, Central Italy

## Abstract

West Nile virus (WNV) and Usutu virus (USUV) are mosquito-borne flaviviruses causing world-wide numerous cases in animals and humans. In Italy, both viruses have been associated with neurological diseases in humans and wild birds. Wild bird rescue centers where support in emergency and care of diseased animals are provided, are potential significant hot spots for avian infection surveillance, as recognized in the Italian Integrate National Surveillance Plan for Arboviruses. Here we report the results of a post-mortem active monitoring study conducted from November 2017 to October 2020 on animals hosted in five wild bird rescue centers of Central Italy. Five hundred seventy-six (n = 576) wild birds were tested by real-time polymerase chain reaction (RT-PCR) for the presence of WNV or USUV RNA fragments. No birds tested positive for USUV RNA (n = 0; 0.00 %). Evidence of WNV RNA (Ct value = 34.36) was found in one bird (n = 1; 0.17 %), an adult little grebe (*Tachybaptus ruficollis* subsp. *ruficollis*), that tested WNV positive on December 2019 and died due to traumatic injuries. The main pathological findings consisted in mild CD3+ lymphocytic tubulo-interstitial nephritis, meningoencephalitis, and cardiomyocytes loss and interstitial oedema of the heart. This study highlights the strategic role of wildlife rescue centers in monitoring both the introduction and circulation of avian emerging zoonotic diseases. Also, the presence of WNV during the cold season evidences the possible role of birds in overwintering mechanisms in the Italian territory and requires further investigations.

## 1. Introduction

The family *Flaviviridae* includes several species within 3 genera: *Flavivirus, Hepacivirus* and *Pestivirus*. Compared to the other two, some species belonging to the genus *Flavivirus* need either mosquito or tick vectors for transmission (Petersen et al., 2005). As vector-borne viruses, their transmission dynamics, geographic spread and re-emergence are greatly influenced by climate change since a warmer climate and changing rainfall patterns may create favorable environments for climate-sensitive vectors (such as mosquitoes and ticks) and pathogens. Although other non-climate factors may play a role, it is common knowledge that climate change is the most important driver influencing the epidemiology and transmission of vector borne pathogens (Patz et al., 2005). Heavy precipitation and extreme heat are likely the principal cause of the increasing number of vector-borne disease outbreaks reported in Europe in the last decades (Moirano et al., 2018; Patz et al., 2005). Most of these outbreaks have been caused by West Nile and Usutu viruses, two species belonging to the *Flavivirus* genus. Included in the Japanese Encephalitis group, they are genetically closely related and share many traits of their transmission cycle which involves birds and mosquitoes. Mammals, including humans, might be infected through mosquito bites and represent dead-end hosts of these infections. When infected, humans usually develop flu-like syndrome although sometimes cases of neuroinvasive disease can occur (Grottola et al., 2017; Pacenti et al., 2019; Santini et al., 2015). In Italy, WNV and USUV infections, sometimes associated with clinical neuroinvasive disease, have been reported in wild birds, horses and humans since late nineties (Cantile et al., 2001; Manarolla et al., 2010; Pacenti et al., 2019; Pautasso et al., 2016; Percivalle et al., 2020, 2017; Scaramozzino et al., 2021; Sinigaglia et al., 2019; Tamba et al., 2011). A national surveillance plan to detect WNV introduction and circulation in the whole country has been implemented by the Directorate-General for animal health and veterinary medicinal products of Animal Health of the Ministry of Health since 2001. Over the years, it has been modified according to the epidemiological evolution of the disease and, in 2017, Usutu virus was included in the plan. Since 2020, an integrated National Plan for Prevention, Surveillance and Response to Arbovirus (PNA) is in place as a five-year plan (Italian Ministry of Health, 2019) and includes, among the others, activities on WNV and USUV. PNA aims to early detect viral circulation in Italy to minimize the risk of human infections. The Italian territory has been classified in three areas according to the epidemiological-environmental conditions and, as a consequence, the transmission risk. *High risk areas* are those where WNV is circulating or has circulated in at least one of the 5 previous years, *low risk areas* are those where WNV has never been or has been rarely reported and where eco-climatic conditions are favorable to viral circulation, *minimum risk areas* are those where WNV has never been reported and where eco-climatic conditions are not suitable to WNV circulation (Italian Ministry of Health, 2019). Whatever the category of the risk area, the surveillance includes monitoring of wild bird mortality. Wild bird rescue centers (WRCs) are scattered in the territory. They are the main providers of medical support in emergency and wild diseased animal care and may represent an important source of information on the WNV and USUV circulation and their virulence characteristics. In this study, carcasses from WRCS of the Central Italy were examined to monitor the circulation of WNV and USUV. To investigate whether wild birds might be an important factor in facilitating the overwintering of USUV and WNV in the study area, bird samples were collected and examined throughout all year.

## 2. Material and Methods

### 2.1. Animals

Carcasses of wild birds (n = 576) collected by 5 rescue centers located in Umbria, Latium, and Tuscany (Central Italy) from November 2017 to October 2020, were used in this study. All birds died spontaneously or were humanely euthanized for clinical conditions compromising animal welfare (e.g., gunshot lesions, head trauma and fractures). The animals were submitted for necropsy at the Department of Veterinary Medicine, University of Perugia, Umbria, Italy. During necropsy, the cadaver condition was scored as to follow: Code 1 (absent autolysis), Code 2 (mild autolysis), Code 3 (moderate autolysis), Code 4 to Code 5 (marked autolysis or corruption, respectively) (Denise Mcaloose, Judy St. Leger, 2018). For animals with Code 1 to 3, tissues collected during the necropsy were submitted to RT-PCR, histopathology and immunohistochemistry analyses to investigate either the presence of WNV or USUV or eventual related lesions; for birds with cadaver condition Code 4 or 5, the collected tissues were submitted only to RT-PCR analyses (Denise Mcaloose, Judy St. Leger, 2018). Ethical approval was not required for this study.

### 2.2. WNV and USUV molecular detection

Samples of heart, liver, spleen, kidney, and brain were collected, stored at -80°C and sent to the WOAH (former OIE) Reference Laboratory for West Nile Fever at the Istituto Zooprofilattico Sperimentale “G. Caporale” (Teramo, Italy) for molecular detection of WNV and USUV RNA by RT-PCR analyses. Tissues (brain and pool of heart, liver, spleen, and kidney) were homogenized in phosphate buffered saline solution. Briefly, after tissue homogenization, RNA extraction was performed by using the MagMAX CORE Nucleic Acid Purification KIT (Applied Biosystem, Termofisher Scientific, Life technologies corporation, TX, USA), following manufacturer’s instructions. The extracted RNA was amplified as described in literature, using a double RT-PCR approach, one aiming for simultaneous detection of WNV lineages 1 and 2 (Del Amo et al., 2013) and the other aiming for detection of all WNV lineages (Vázquez et al., 2016). For USUV, the RT-PCR was performed as previously described by Cavrini et al., 2011.

### 2.3. Histopathology and Immunohistochemistry (IHC)

For animals with Code 1 to 3 routine histological examination was performed on FFPE tissues collected during necropsy. For the immunohistochemistry, 3 μm sections were obtained from FFPE tissues; the sections were deparaffinized and rehydrated in alcohols. Endogenous peroxidases were blocked by 3% H_2_O_2_ in methanol incubation for 10 minutes and antigen was retrieved by proteinase K digestion for 10 minutes at 37°C. A goat serum block was applied before the incubation with the primary antibody. For primary incubation, anti-WNV serum (FLI, Germany) was used with 1:1700 dilution. As secondary antibody, Bright vision 1 step detection system anti-rabbit HRP was used, with the aminoethyl carbazole (AEC) as substrate. For USUV, IHC was performed as previously described (Giglia et al., 2021).

Primary polyclonal Rabbit Anti-CD3 antibody was used at a dilution of 1:200 (A0452, Dako) to mark T-cells as common player of adaptive immunity to intracellular agents in inflammatory infiltrates (Kapczynski and Segovia, 2020). As secondary antibody, Mouse Envision HRP kit was used (Ab93697; Abcam) with the aminoethyl carbazole (AEC) as substrate.

### 2.4. Geographical distribution analysis

To map geographical distribution of the rescued animals, collection sites were registered, and coordinates recorded. If the exact collection site was not available, data of the rescue center were used. To visualize the geographical distribution of the collected animals and the location of identified cases, the open-source Quantum Geographic Information System (QGIS®) (v. 3.16.10) was used. The geographical distribution of sampling sites and positive case location is shown in Figure 1.

**Figure 1.**
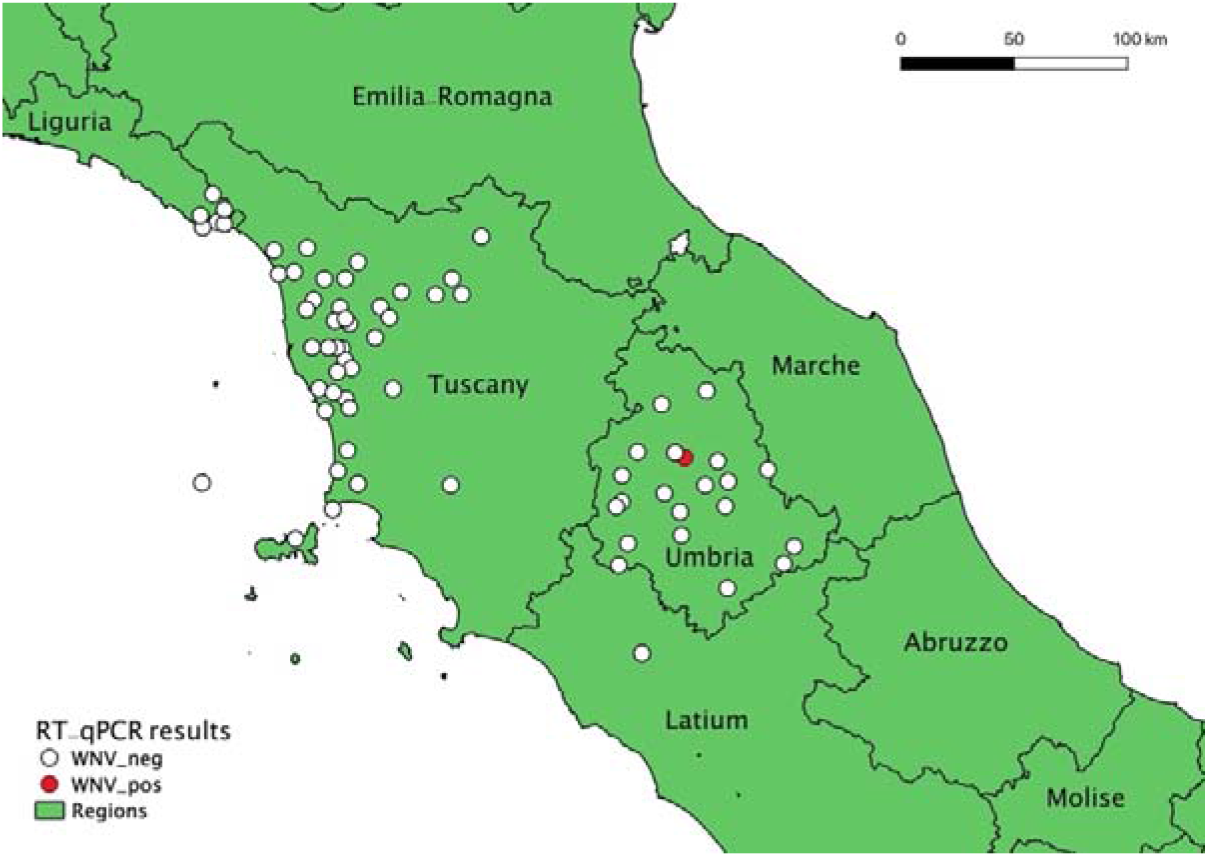
Geographical distribution of sampling sites and positive case location (QGIS map). The map shows the sites of wild birds sampling (white dots) distributed in Umbria, Lazio, Tuscany, and the location of the WNV-positive little grebe (*Tachybaptus ruficollis* subsp. *ruficollis*) (red dot).

## 3. Results

One out of 576 (0.17%) wild birds tested for the presence of WNV and USUV RNA was found positive for WNV (Ct value: 34.36), while none of the wild birds tested was positive for USUV (Table 1). The only WNV positive bird was a little grebe (*Tachybaptus ruficollis* subsp. *ruficollis*) belonging to the Order *Podicipediformes*, Family *Podicipedidae*. The bird was found on the street side in the Umbria region at the end of December and admitted to the Veterinary Teaching Hospital of the Department of Veterinary Medicine of the University of Perugia (Italy). It showed depression and difficulties in the movements, partially justified by the presence of a complete luxation of the 3^rd^ phalanx of the 3^rd^ digit, confirmed by radiography. The bird died spontaneously during the day after admission. At necropsy, a mild hepatomegaly and multiorgan congestion were observed. At microscopy, the heart showed mild multifocal loss of cardiomyocytes and interstitial oedema. In kidneys tubulo-interstitial nephritis and tubular necrosis were observed, while in the brain, scattered lymphocytes were seen in the meninges and perivascular neuroparenchyma. At immunohistochemistry, leucocyte infiltration consisted in CD3-positive lymphocytes. WNV and USUV antigen were not detected.

**Table 1.**
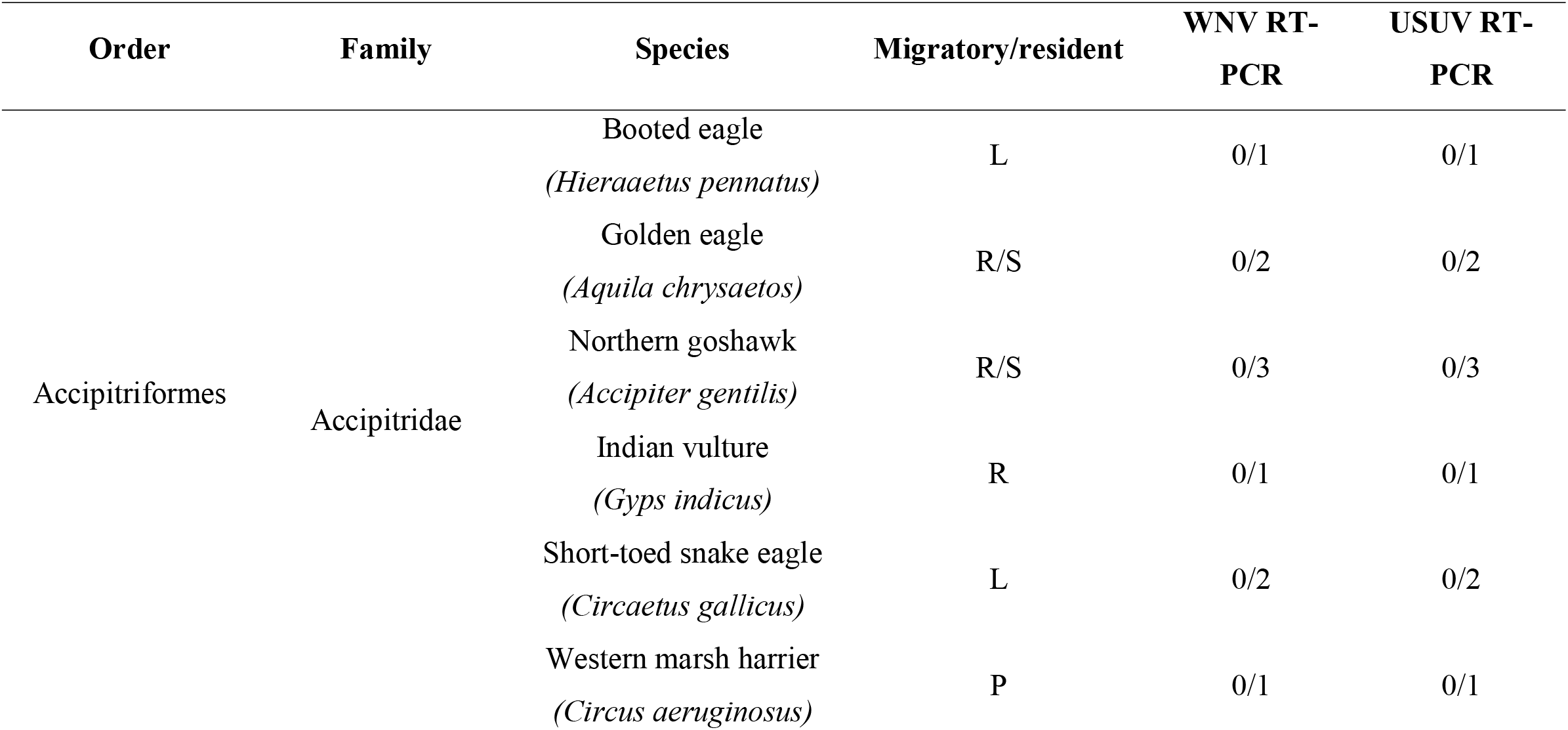
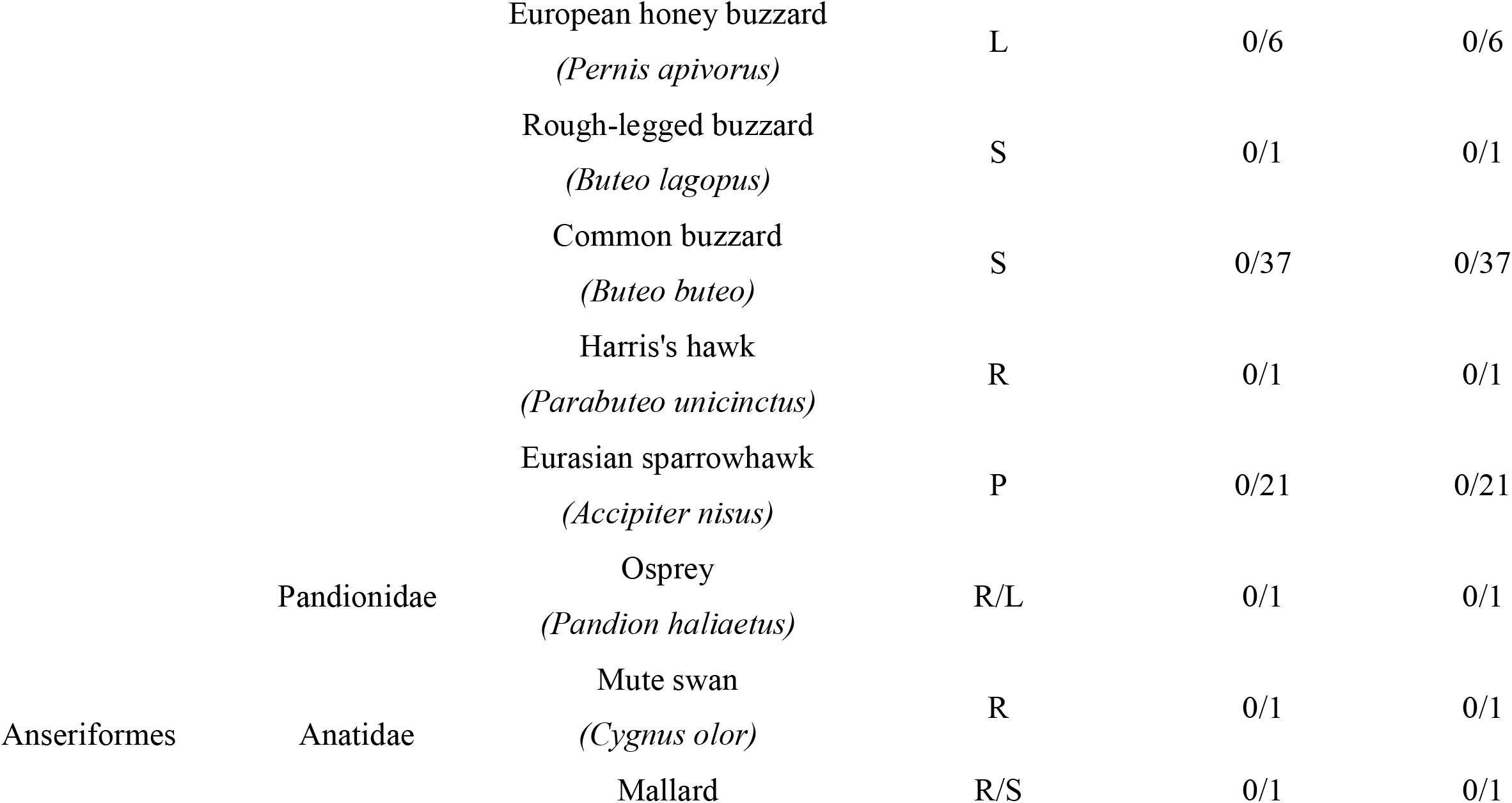
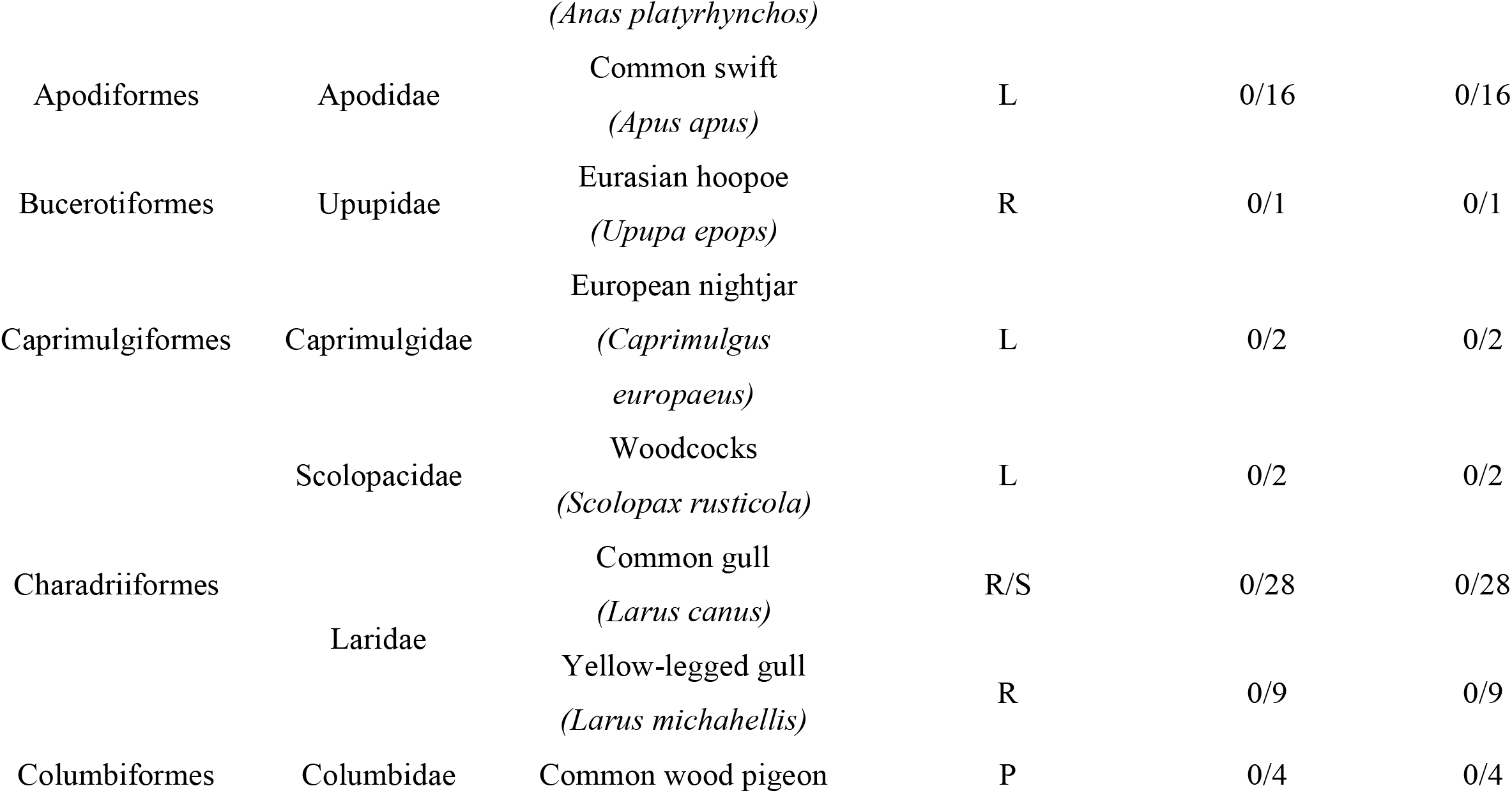
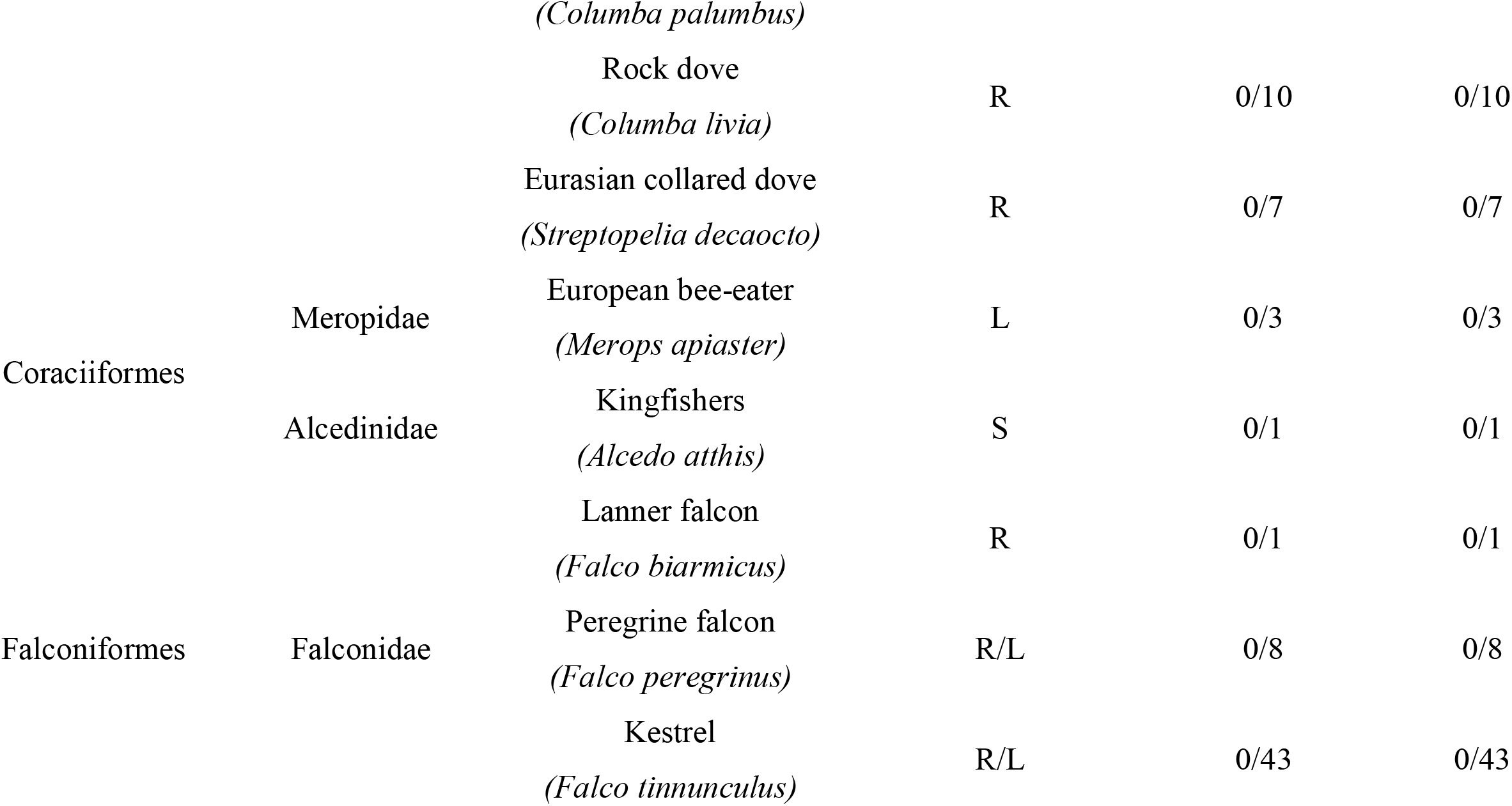
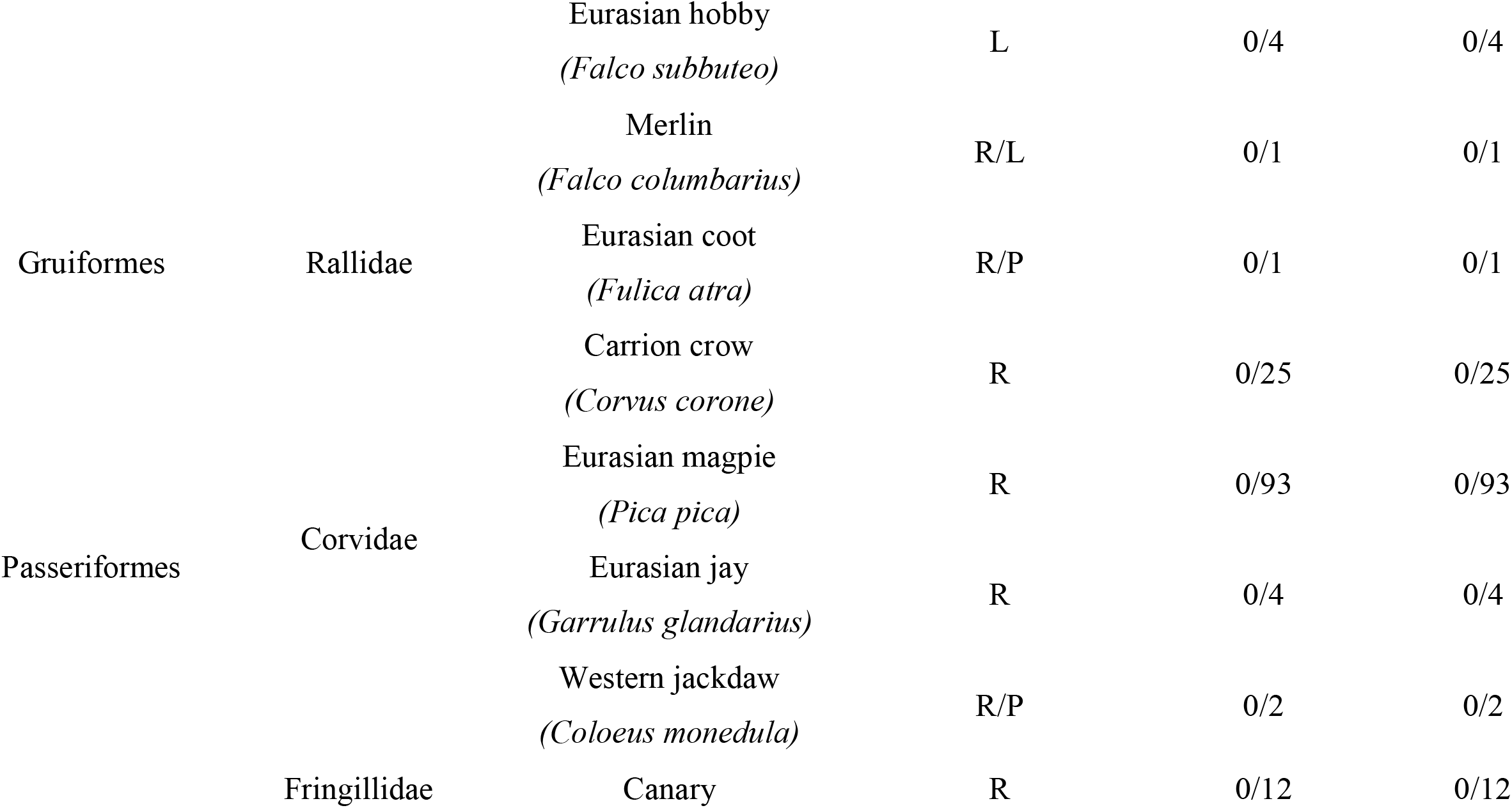
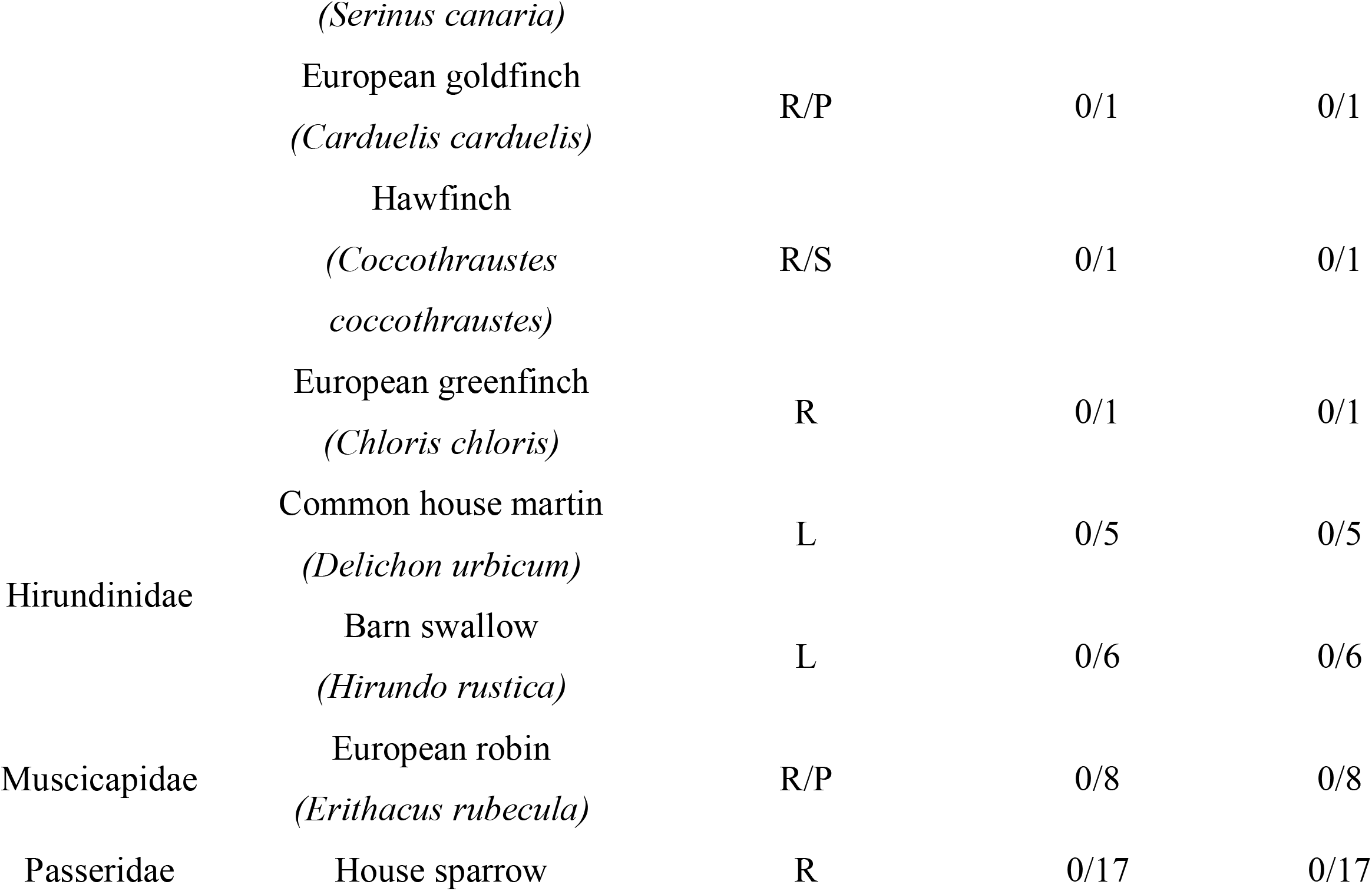
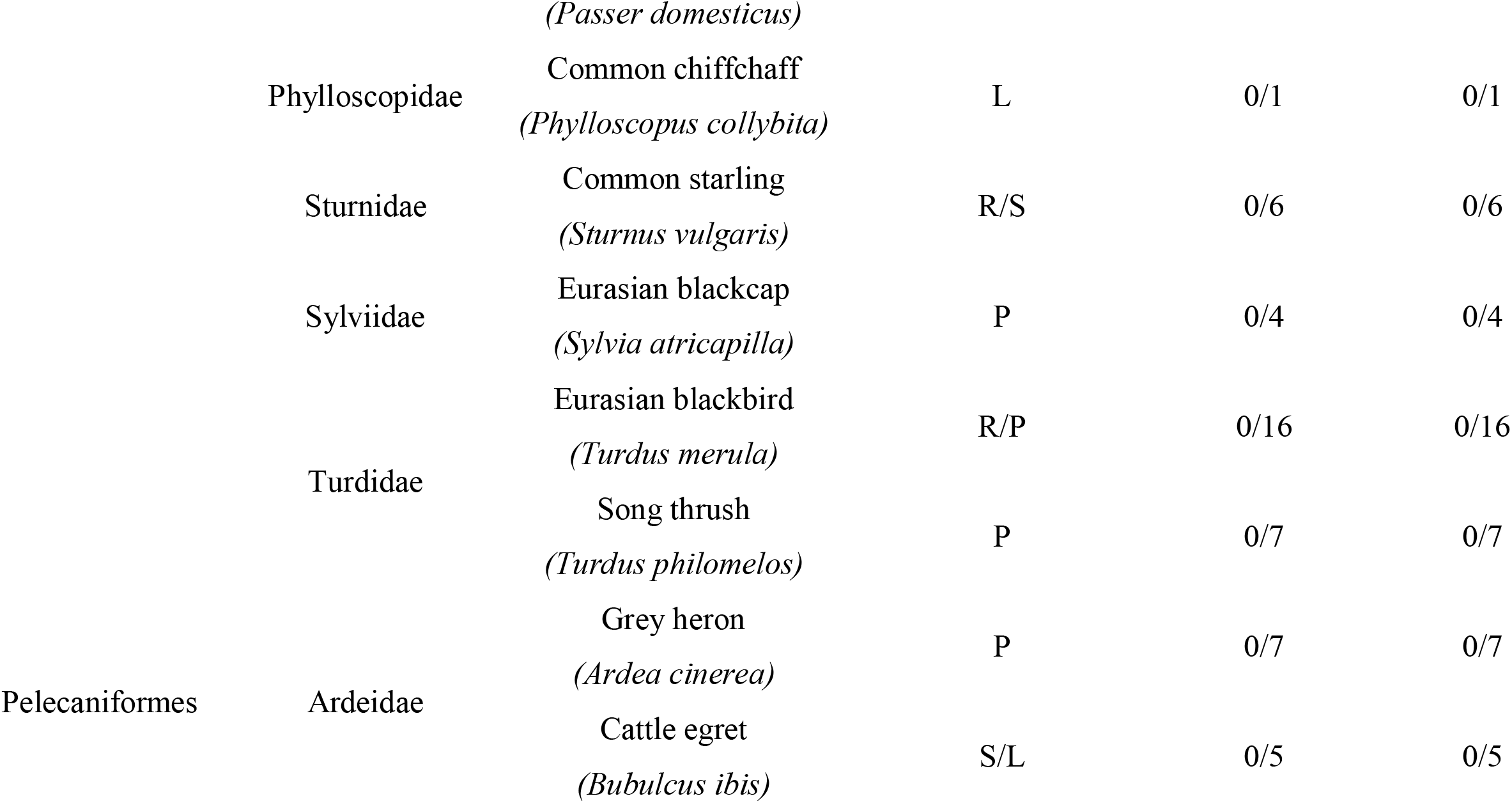
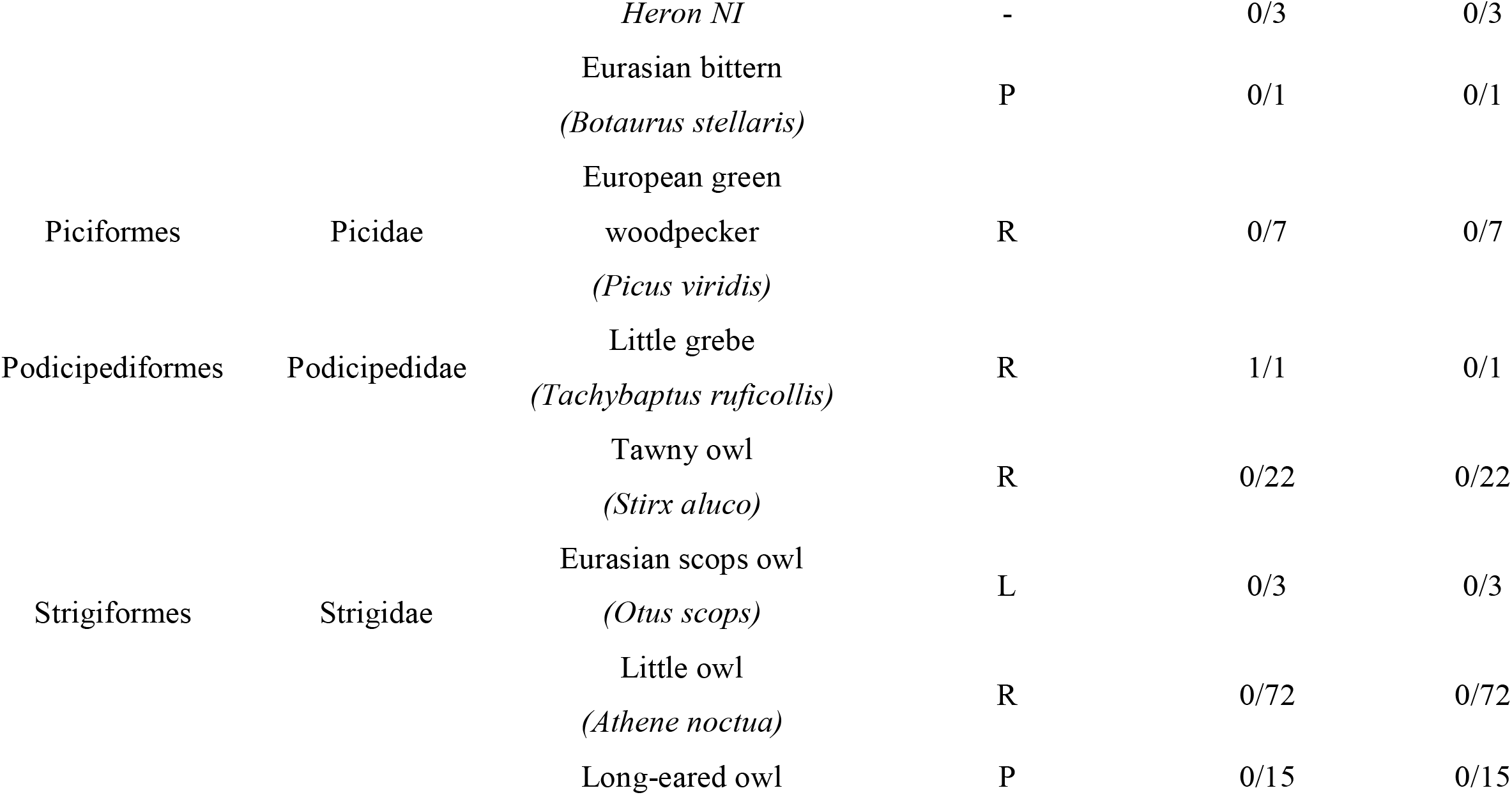
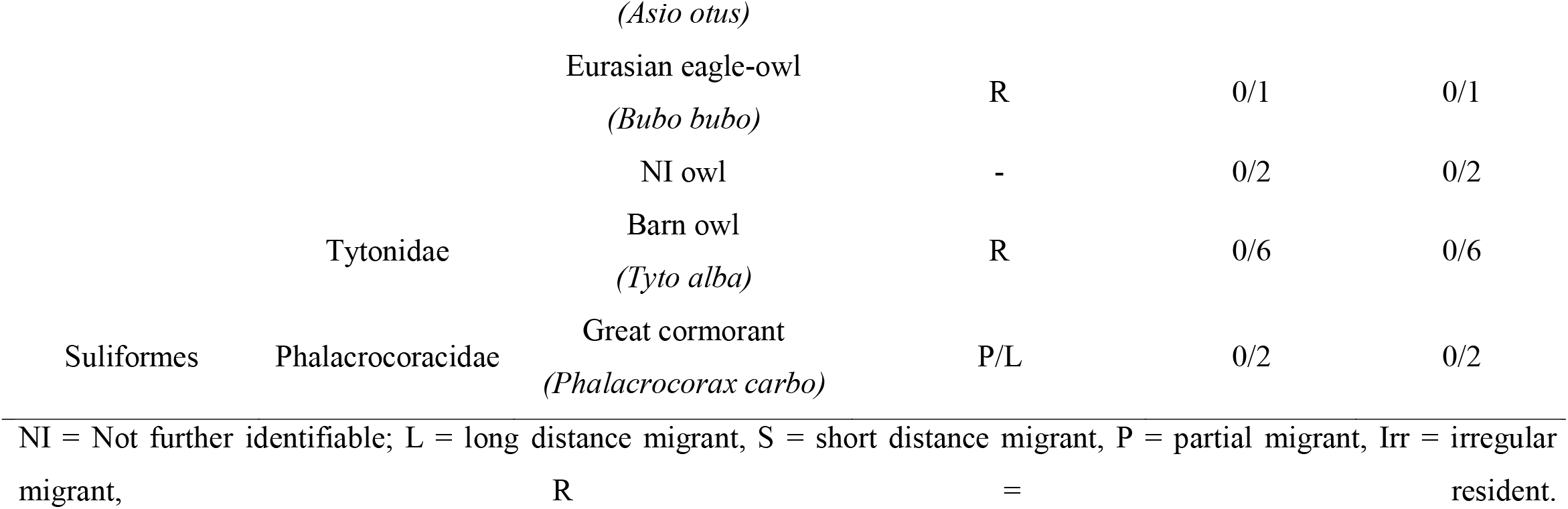
Summary of examined avian species, migratory behavior and WNV and USUV RT-PCR results.

## 4. Discussion

Epornitic mosquito-borne zoonotic flaviviruses associated with cases in animals and humans have significantly increased their impact in public health in the last decades (Benzarti et al., 2019; Vilibic-Cavlek et al., 2020). Their continuous circulation throughout Europe have raised the attention and supported the development of National Surveillance Systems to early detect virus circulation and limit its spread (Pacenti et al., 2019). For mosquito-borne zoonotic flavivirus circulation, birds, being the amplifying hosts, are the major target in surveillance strategies (Hubálek et al., 2014; Vilibic-Cavlek et al., 2020).

This study reports a post-mortem monitoring on wild birds admitted in WRCs of Central Italy between November 2017 and October 2020. According to the WNV risk classification areas as established in the PNA (Italian Ministry of Health, 2019), the WRCs involved in this study were located in the low (Umbria) and in the high risk areas (Latium and Tuscany).

WNV lineage 2 was detected a little grebe (*Tachybaptus ruficollis* subsp. *Ruficollis;* Order *Podicipediformes*, Family *Podicipedidae*). Little grebe is in the IUCN Red List of Threatened Species (2019). It inhabits small and shallow wetlands and for this reason it is considered resident in many European countries. Being a resident species implies that it has probably been infected within the rescue area. In other words, it implies that the infection likely occurred in Umbria. As far as we know, this was the first evidence of WNV circulation in Umbria, an area classified as low risk for WNV circulation. However, looking further, finding a WNV positive bird in Umbria was somewhat expected as USUV, which shares same ecological niche of WNV, has been repeatedly detected in this region (). So, conditions for WNV to establish in the area existed. Conversely, finding a WNV positivity at the end of December, when vectors are not flying, was rather surprising. In this case, the histological lesions were mild, antigen couldn’t be detected, and the Ct values were quite high (34.36 cycles). These results if on the one hand exclude the hypothesis of a recent infection, on the other hand, they might instead suggest the presence of a persistent infection (Reisen et al., 2013). Persistent infection has been defined as the detection of virus in host tissues after viremia has receded (Wheeler et al., 2012). The persistent viral load in organs of birds, and in particular in those belonging to prey species like the little grebe, might sustain the WNV transmission to predators also months after mosquito season (Mencattelli et al. 2022). In our case this implies the possibility of WNV to overwintering in Umbria through the bird-to-bird transmission.

The little grebe was the only bird out of 576 (0.17%) found positive to WNV RT-qPCR. None of the birds tested positive to USUV RT-qPCR.

The lack of WNV and USUV infection cases in high risk areas (Latium and Tuscany) also during low vector activity periods, additionally support the idea previously suggested in literature that contrarily to vectors, birds have a minor role in the overwintering of these viruses (Rudolf et al., 2017). Alternatively, the high number of negative results and, in particular, of those found in the high-risk areas might be indicative of the low virulence of most of the WNV and USUV strains circulating in these regions.

Our results also demonstrate that even if not being ideal to be used all alone in a surveillance program because they rescue only diseased or injured animals, WRCs can instead be very useful to the National Surveillance plan as they can improve its sensitivity, expand the period of surveillance by including the winter months and provide important information on the virulence of the circulating strains. As a final point, considering that the little grebe is a threatened species, we cannot underestimate that WNV might play a role in this species loss.

## 5. Conclusions

For the first time, a case of WNV infection in Umbria (Central Italy), a region currently classified as low-risk area for WNV, was detected during a low vector activity season (winter) This result suggests that the active monitoring performed through the National Plan of Surveillance on wild birds from wild bird rescue centers can help to better detect the introduction and circulation of WNV. The lack of detection of USUV circulation among mosquitos and birds needs to be monitored to keep assuring early detection in humans and animals. To assure WNV circulation and infections of animals and humans are kept under control, additional systematic epidemiological plans of active surveillance on animals received at rescue centers in Italy are highly recommended.

